# Internal selective attention is delayed by competition between endogenous and exogenous factors

**DOI:** 10.1101/2022.07.05.498906

**Authors:** Edward F. Ester, Asal Nouri

## Abstract

External selective attention is mediated by competition between endogenous (goal-driven) and exogenous (stimulus-driven) factors, with the balance of competition determining which stimuli are selected. Occasionally, exogenous factors “win” this competition and drive the selection of task-irrelevant stimuli. Endogenous and exogenous selection mechanisms may also compete to control the selection of internal representations (e.g., those stored in working memory), but how this competition manifests and whether it is resolved in the same way as external attention is unknown. Here, we leveraged the high temporal resolution of human EEG to determine how competition between endogenous and exogenous factors influences the selection of internal representations. Unlike external attention, competition between endogenous and exogenous factors did not prompt the selection of task-irrelevant working memory content. Instead, it simply delayed the endogenous selection of task-relevant working memory content by several hundred milliseconds. Thus, competition between endogenous and exogenous factors influences internal selective attention, but in a different way than external selective attention.

## Significance Statement

Selective attention can be allocated to external stimuli based on goal-driven or stimulus-driven factors. These factors often compete, and the balance of competition determines where we pay attention. A similar type of competition may govern the selection of internal representations, i.e., those stored in working memory. We tested whether (a) goal-and stimulus-driven factors compete to control the selection of information stored in working memory, and (b) whether this competition results in the selection of irrelevant memory content akin to that seen during the selection of external stimuli. We find clear evidence for a competition between goal- and stimulus-driven factors during the selection of working memory content, but no evidence for the selection of task-irrelevant memoranda. Thus, different mechanisms may govern the goal- and stimulus-driven selection of external sensory inputs and internal working memory representations.

Efficient behavior requires rapid comparison of sensory inputs with internal representations of goal states and motor affordances. Many of these comparisons take place in working memory (WM), a capacity- and duration-limited system that forms a temporal bridge between fleeting sensory phenomena and possible actions (D’Esposito & Postle, 2015; van Ede & Nobre, 2023). Capacity limits in WM necessitate the existence of external selection mechanisms that gate access to this system (i.e., input gating), while rapidly changing environmental circumstances necessitate the existence of internal selection mechanisms that prioritize behaviorally relevant subsets of information stored in WM for action (i.e., output gating). Whether similar mechanisms mediate the selection of internal and external information is hotly debated (Chun et al., 2011; Chatham & Badre, 2015; Rac-Lubashevsky & Frank, 2021).

External sensory inputs can be selected based on behavioral goals (i.e., endogenous selection) or stimulus properties (i.e., exogenous selection), with selection ultimately determined by the balance of competition between these factors. For example, stimulus factors can trigger the selection of task-irrelevant information (Theeuwes, 1992; Folk et al., 1992; Anderson et al., 2011), disrupting top-down searches for task-relevant stimuli (Wolfe, 2020). These disruptions are frequently accompanied by concurrent shifts in cortical and subcortical spatial priority maps that mediate eye movements and endogenous shifts of covert spatial attention (e.g., Bisley & Goldberg, 2013; Sprague & Serences, 2013; Luck et al., 2020).

Endogenous and exogenous factors may also compete to control the selection of internal representations, for example, those stored in WM (e.g., van Ede et al., 2020). However, little is known about how this competition influences memory performance and is resolved. One obvious possibility is that competition results in the exogenous selection of task-irrelevant information like that seen in external attention. For example, an external stimulus might trigger the exogenous selection of stimulus-matching WM content (i.e., the converse of WM-guided selection, where external attention is oriented to task-irrelevant stimuli that incidentally match attributes of stimuli stored in WM; Olivers et al., 2011). Alternately, competition between endogenous and exogenous factors could produce a general slowing or delay in the selection of task-relevant memory content without prompting the exogenous selection of task-irrelevant memory content. This may explain a recent finding documenting delays in oculomotor biases to the locations of items stored in WM when experimental factors place endogenous and exogenous selection mechanisms in conflict (van Ede et al., 2020).

To test these possibilities, we recorded EEG while participants performed a retrospectively cued WM task typically used to study internal attention (e.g., Griffin & Nobre, 2003; Landman et al., 2003). In different experimental blocks, a cue presented during WM storage indicated which of two memorized positions would be probed for recall (pro-cue trials) or which position would not be probed for recall (anti-cue trials). We reasoned that the anti-cue manipulation would create a state of conflict between endogenous and exogenous selection mechanisms, a point supported by studies documenting visual search costs when participants are cued to the identity of an upcoming distractor (e.g., Moher & Egeth, 2012; van Ede et al., 2020). We then examined how informative pro- and anti-cues influenced EEG signals that track covert shifts of spatial attention with high temporal precision. Across multiple analyses, we found no evidence for shifts of attention toward cue-matching but task-irrelevant memory representations during the anti-cue task. Instead, we observed a significant delay in the selection of task-relevant WM content during the anti-cue relative to the pro-cue task. Control analyses demonstrated that this result could not be explained by weaknesses in our experimental design or idiosyncrasies in our analytic approach. Thus, we argue that unlike external attention – where competition between endogenous and exogenous selection mechanisms can produce the selection of task-irrelevant information – competition between endogenous and endogenous internal selection mechanisms does not produce an exogenous selection of task-irrelevant information and is instead resolved in a fundamentally different way.

## Methods

### Participants

42 human volunteers (both sexes) participated in a single 2.5-hour testing session. Participants were recruited from the Florida Atlantic University community via campus advertisements and remunerated at $15/h in Amazon.com gift cards. All participants gave both written and oral informed consent in compliance with procedures established by the local institutional review board, and all participants self-reported normal or corrected-to-normal visual acuity. Two participants voluntarily withdrew from the study prior to completing both cue conditions (i.e., pro-cue vs. anti-cue); data from these participants were excluded from final analyses. Thus, the data reported here reflect the remaining 40 participants.

### Testing Environment

Participants were seated in a dimly-lit and acoustically shielded recording chamber for the duration of the experiment. Stimuli were generated in MATLAB and rendered on a 17’’ Dell CRT monitor cycling at 85 Hz (1024 x 768 pixel resolution) via PsychToolbox3 software extensions. Participants were seated approximately 65 cm from the display (head position was unconstrained). To combat fatigue and/or boredom, participants were offered short breaks at the end of each experimental block.

### Spatial Retrocue Task

A task schematic is shown in Figure 1A. Each trial began with the presentation of an encoding display lasting 500 ms. The encoding display contained two colored circles (blue and red; subtending 1.75° visual angle from a viewing distance of 65 cm) rendered at of eight polar locations (22.5° to 337.5° in 45° increments) along the perimeter of an imaginary circle (radius 7.5° visual angle) centered on a circular fixation point (subtending 0.25°) rendered in the middle of the display. The locations of the two discs were counterbalanced across each task (i.e., pro-cue vs. anti-cue), though not necessarily within an experimental block. Participants were instructed to maintain fixation and refrain from blinking for the duration of each trial.

**Figure 1.**
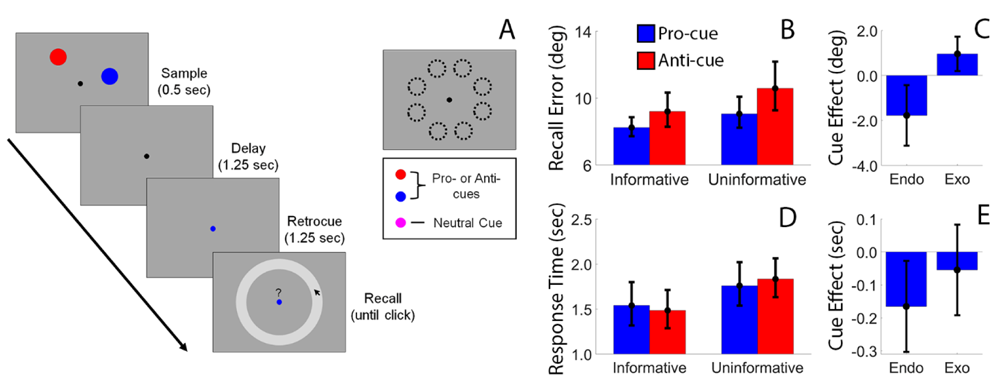
Memory Task and Performance. (A) Participants remembered the locations of two discs over a blank delay. Each disc could appear at one of eight positions along the perimeter of an imaginary circle centered at fixation (upper right panel). (B) Effects of cue type (informative, uninformative) and task type (pro-cue, anti-cue) on average absolute recall errors. (C) We estimated the effects of exogenous factors on recall performance by computing the difference between informative pro-cue trials (i.e., where endogenous and exogenous factors are aligned) and informative anti-cue trials (i.e., where endogenous and exogenous factors are opposed). We estimated the effects of endogenous factors on recall performance by computing the difference between informative pro-cue trials and uninformative pro-cue trials minus the estimated effect of exogenous factors (see text for specifics). Identical analyses were also applied to participants response times (D, E). Error bars depict the 95% confidence interval of the mean.

The sample display was followed by a 1.25 sec blank display and a 1.25 sec retrocue display. Retrocues were defined by a change in the color of the fixation point; during informative trials the fixation point changed colors from black to either blue or red (i.e., matching the color of a remembered disc). During neutral trials, the fixation point changed colors from black to purple (the “average” of blue and red). At the end of the trial, a response display containing a mouse cursor, “?” symbol, and a probe circle appeared. Participants were required to report the location of a remembered disc by clicking along the perimeter of the probe circle (during neutral trials, the fixation point changed colors from purple to either blue or red during the response display, indicating which item should be reported). Participants were instructed to prioritize accuracy over speed, and no response deadline was imposed. The trial terminated as soon as the participant clicked on a location. Sequential trials were followed by a 1.5-2.5 sec blank period (randomly and independently selected from a uniform distribution after each trial).

During the first half of the experiment (e.g., experimental blocks 1-8), each participant was assigned to the pro-cue or anti-cue task. In the pro-cue task, participants were instructed that during informative trials they would be required to click on the location of the disc matching the retrocue color. Conversely, in the anti-cue task participants were instructed that during informative trials they would be required to click on the location of the disc that did not match the retrocue color. Participants completed eight blocks of 56 trials in the both the pro-cue and anti-cue tasks. Task order (i.e., eight blocks of the pro-cue task followed by eight blocks of the anti-cue task or vice versa) was counterbalanced across participants.

### EEG Acquisition and Preprocessing

Continuous EEG was recorded from 63 uniformly distributed scalp electrodes using a BrainProducts “actiCHamp” system. The horizontal and vertical electrooculogram (EOG) were recorded from bipolar electrode montages placed over the left and right canthi and above and below the right eye, respectively. Live EEG and EOG recordings were referenced to a 64^th^ electrode placed over the right mastoid and digitized at 1 kHz. All data were later re-referenced to the algebraic mean of the left- and right mastoids, with 10-20 site TP9 serving as the left mastoid reference.

Data preprocessing was carried out via EEGLAB software extensions (Delorme & Makeig, 2004) and custom software. Data preprocessing steps included the following, in order: (1) resampling (from 1 kHz to 250 Hz), (2) bandpass filtering (1 to 50 Hz; zero-phase forward- and reverse finite impulse response filters as implemented by EEGLAB), (3) epoching from -1.0 to +5.0 sec relative to the start of each trial, (4) identification, removal, and interpolation of noisy electrodes via EEGLAB software extensions, and (5) identification and removal of oculomotor artifacts via independent components analysis as implemented by EEGLAB. After preprocessing, location decoding analyses focused exclusively on the following 10-20 occipitoparietal electrodes: P7, P5, P3, Pz, P2, P4, P6, P8, PO7, PO3, POz, PO2, PO4, PO8, O1, O2, Oz.

### Data Cleanup

Prior to analyzing participants’ behavioral or EEG data, we excluded all trials where the participant responded with a latency of < 0.4 sec (we attributed these trials to accidental mouse clicks following the onset of the probe display rather than a deliberative recall of a specific stimulus position) and more than 3 standard deviations above the average response time across all experimental conditions. This resulted in an average (±1 S.E.M.) loss of 14.43 ±0.93 trials (or 1.67% ± 0.11% of trials) across participants but had no qualitative effect on any of the findings reported here.

### Decoding Spatial Positions from Posterior Alpha-Band EEG Signals

Location decoding was based on the multivariate distance between EEG activity patterns associated with memory for specific positions. This approach is an extension of earlier parametric decoding methods (Wolff et al., 2017) designed for use in circular feature spaces. We extracted spatiotemporal patterns of alpha-band activity (8-13 Hz) from 17 occipitoparietal electrode sites (see *EEG Acquisition and Preprocessing* above). The raw timeseries at each electrode was bandpass filtered from 8-13 Hz (zero-phase forward-and-reverse filters as implemented by EEGLAB software), yielding a real-valued signal f(t). The analytic representation of f(t) was obtained via Hilbert transformation:

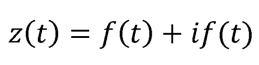

where *i* is the imaginary operator and *if(t)* = *A(t)e*^*iφ(t)*^. Alpha power was computed by extracting and squaring the instantaneous amplitude A(t) of the analytic signal z(t).

Location decoding performance was computed separately for each disc (i.e., blue vs. red), trial type (i.e., informative vs. neutral) and each task (i.e., pro-cue vs. anti-cue) on a timepoint-by-timepoint basis. In the first phase of the analysis, we sorted data from each condition into 5 unique training and test data sets using stratified sampling while ensuring that each training set was balanced across remembered positions (i.e., we ensured that each training data set contained an equal number of observations where the location of the remembered stimulus was at 22.5°, 67.5°, etc.). We circularly shifted the data in each training and test data set to a common center (0°, by convention) and computed trial-averaged patterns of responses associated with memory for each disc position in each training data set. Next, we computed the Mahalanobis distance between trial-wise activation patterns in each test data set with position-specific activation patterns in the corresponding test data set, yielding a location-wise set of distance estimates. If scalp activation patterns contain information about remembered positions then distance estimates should be smallest when comparing patterns associated with memory for identical positions in the training and test data set and largest when comparing opposite positions (i.e., those ±180° apart), yielding an inverted gaussian-shaped function. Trial-wise distance functions were averaged and sign-reversed for interpretability. Decoding performance was estimated by convolving timepoint-wise distance functions with a cosine function, yielding a metric where chance decoding performance is equal to 0. Decoding results from each training- and test-data set pair were averaged (thus ensuring the internal reliability of our approach), yielding a single decoding estimate per participant, timepoint, and task condition. To verify that our findings are not contingent on the specific type of decoding analysis used, we repeated the aforementioned analysis via eight-way support-vector-machine (SVM) based classification using a “one-versus-all” motif (see Results).

### Cross-correlation Analyses

Temporal differences in task-relevant location decoding performance were estimated via cross-correlation analyses. For neutral trials, we extracted task-relevant decoding performance during pro- and anti-cue blocks over a period spanning 0.0 to 1.0 seconds following the onset of the probe display (i.e., when an informative cue instructed participants which disc to recall). We computed correlation coefficients between pro- and anti-cue decoding time series while systematically shifting the pro-cue time series from -1.0 to +1.0 sec relative to the anti-cue decoding time series (e.g., Figure 3D, blue line). We compared these correlation coefficients to a distribution of correlation coefficients computed under the null hypothesis (i.e., no systematic difference in pro- and anti-cue decoding time series) by repeating the same analysis 10,000 times while randomizing the decoding condition labels (i.e., pro- vs. anti-cue) for each participant. An identical analysis was performed on task-relevant pro- and anti-cue decoding task performance from 0.0 to 1.75 during informative trials. We deliberately selected a longer temporal interval for analysis during informative trials as we expected increases in pro- and anti-cue decoding performance to begin during the retrocue period and persist into the ensuing response period.

**Figure 2.**
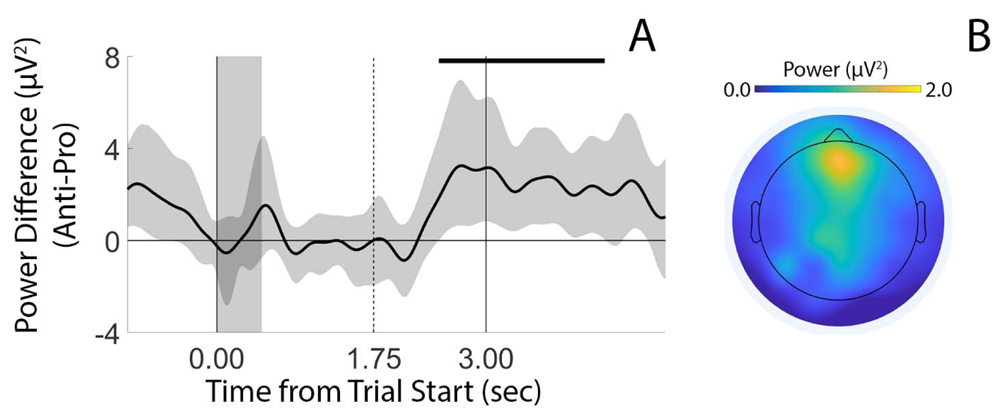
Frontal Theta Power is Greater During the Anti- vs. Pro-Cue Task, Reflecting a Greater Need for Cognitive Control. (A) Time-resolved differences in pro- and anti-cue theta power computed from frontal electrode sites. Theta power estimates were larger during the antivs. pro-cue task beginning approximately 600 ms after cue onset. Shaded regions depict the 95% confidence interval of the mean. Vertical solid lines at times 0.00 and 3.00 depict the onset of the sample and recall displays, respectively; the vertical dashed line at time 1.75 depicts the onset of an informative retrocue. The horizontal black bar at the top of the plot marks periods where the difference between anti- and pro-cue theta power was significantly greater than zero (cluster-based permutation tests; see Methods). (B) Difference in theta-power (4-7 Hz) scalp topography during the pro- and anti-cue tasks. Pro- and anti-cue theta power estimates were averaged over a period spanning 2.5-3.0 sec after trial start (i.e., 750-1250 ms after cue onset). Electrode-wise power estimates during the pro-cue task were subtracted from corresponding estimates during the anti-cue task, i.e., larger values indicate higher theta power during the antivs. pro-cue task.

**Figure 3.**
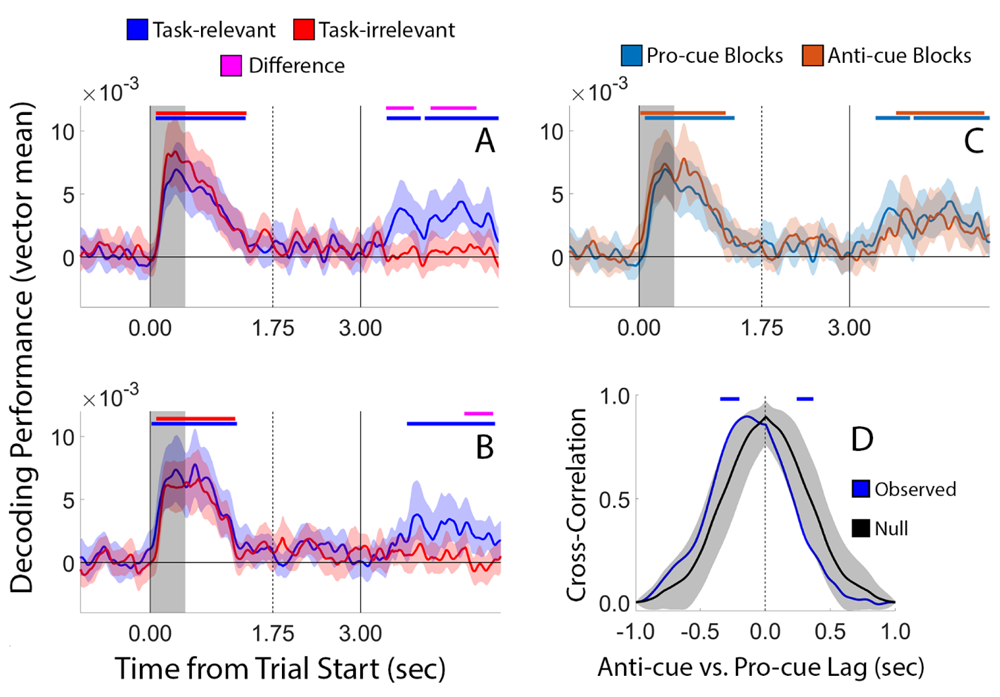
Location Decoding Performance During Neutral Trials. (A, B) Decoding performance for task-relevant and task-irrelevant locations during pro-cue and anti-cue blocks, respectively. (C) Overlay of task-relevant location decoding performance for pro-cue and anticue blocks (i.e., the blue lines in panels A and B). Solid vertical lines at time 0.00 and 3.00 depict the onset of the sample and probe displays, respectively. The dashed vertical line at time 1.75 depicts the onset of the (uninformative) retrocue. Gray shaded region spanning 0.00-0.50 marks the duration of the sample display. Horizontal bars at the top of each plot mark intervals where decoding performance was significantly greater than zero (nonparametric cluster-based randomization test; see Methods) or intervals where decoding performance for one location was significantly greater than decoding performance for the other location. Shaded regions around each line depict bootstrapped confidence intervals of the mean. (D) Cross-correlation analysis showing a significant delay in the onset of above-chance probe-locked task-relevant decoding performance during the anti- vs. pro-cue task. The null distribution was obtained by repeating the cross-correlation analysis 10,000 times while randomizing participant-level condition labels (i.e., randomly switching the pro- and anti-cue labels). Horizontal bars at the top of the plot depict intervals where the observed cross-correlation coefficient was significantly greater than that expected by chance.

### Quantifying Frontal Theta Power

Analyses of frontal theta power focused on informative trials from the pro- and anti-cue tasks. The raw timeseries at each scalp electrode was bandpass filtered from 4-7 Hz (zero-phase forward-and-reverse filters as implemented by EEGLAB software), yielding a real-valued signal f(t). The analytic representation of this signal was obtained via Hilbert transformation, and theta power was computed by extracting and squaring the instantaneous amplitude A(t) of the analytic signal z(t). Topographic maps of theta power during the pro- and anti-cue tasks were obtained by averaging power estimates over trials and a temporal window spanning 2.5 to 3.0 sec following the start of each informative trial (i.e., 750 to 1000 ms after cue onset). Based on these maps, we limited further analyses to power estimates measured at four frontal electrode sites: AFz, Fz, F1, and F2. Data from these electrodes were used to compute time-resolved estimates of theta power during the pro- and anti-cue tasks and task differences in theta power. In a final analysis, we extracted and computed trial-wise estimates of theta power during the anti-cue task (using the same electrodes and temporal window described in the previous paragraph). We sorted participants’ anti-cue EEG data into low- and high-theta power groups after applying a media split to theta power estimates, then decoded the location of the cue-matching but task-irrelevant item within each group. This allowed us to test whether evidence for exogenous selection of and/or “refreshing” of cue-matching memory traces was more likely to occur on trials with low- vs. high theta power.

### N2pc Analyses

The N2pc is a negative-going waveform defined by a greater negativity over occipitoparietal electrode sites contralateral vs. ipsilateral to the hemifield containing a visual target (Luck & Hillyard, 1994). We used the N2pc to track covert spatial selection of the cue-matching and task-relevant position during pro-cue blocks and the cue-matching but task-irrelevant stimulus during anti-cue blocks. To control for sensory imbalances across visual hemifields, we restricted our analysis to trials where the task-relevant and task-irrelevant discs appeared in opposite visual hemifields. We estimated voltages over occipitoparietal electrode site pairs O1/2, PO3/4, and PO7/8 during trials when the task-relevant stimulus was in the left vs. right visual field, then sorted trial-wise voltage estimates by the hemifield containing the task-relevant target, i.e., contralateral vs. ipsilateral. We defined the N2pc as the average difference in voltage across contralateral and ipsilateral electrode sites over a period spanning 200-300 ms.

### Inverted Encoding Model

To verify the generality of our findings across analytic approaches, we reconstructed position-specific WM representations from spatiotemporal patterns of alpha-band activity using an inverted encoding model. Our approach was conceptually and quantitatively identical to that used in earlier studies (e.g., Sprague et al., 2016; Ester et al., 2018; Nouri & Ester, 2020). We modeled alpha power at each scalp electrode as a weighted sum of eight location-selective channels, each with an idealized tuning curve (a half-wave rectified cosine function raised to the 8^th^ power, with the maximum response of each function normalized to 1). The predicted responses of each channel during each trial were arranged in a *k* channels by *n* trials design matrix C. The relationship between the EEG data and the predicted responses in *C* is given by a general linear model of the form:

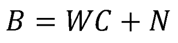

where *B* is an *m* electrode by *n* trial training data matrix, W is an *m* electrode by *k* channel weight matrix, and *N* is a matrix of residuals (i.e., noise).

To estimate *W*, we constructed a training data set containing an equal number of trials for each stimulus position (i.e., 22.5-337.5° in 45° increments). We identified the location φ with the fewest *r* repetitions and constructed a training dataset *Btrn* (*m* electrodes by *n* trials) and weight matrix *Ctrn* (*n* trials by *k* channels) by randomly selecting (without replacement) 1 to *r* trials for each of the eight possible stimulus positions. The training dataset was used to compute a weight for each channel *Ci* using ordinary least-squares estimation:

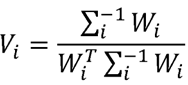

where *T* and -1 are the matrix transpose and inversion operations, respectively. Σ*i* is the regularized noise covariance matrix for each channel *i*, estimated as:

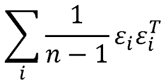

where *n* is the number of training trials and ε*i* is a matrix of residuals:

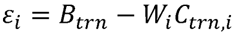

Estimates of ε*i* were obtained by regularization-based shrinkage using an analytically determined shrinkage observation. In this way, an optimal spatial filter *vi* was estimated for each channel *Ci*, yielding an *m* electrode by *k* filter matrix *V*.

Next, we constructed a test dataset *Btst* (*m* electrodes by *n* trials) containing data from all trials not included in the training data set and estimated trial-by-trial channel responses *Ctst* (*k* channels by *n* trials):

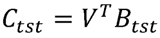

Trial-wise channel response estimates were interpolated to 360°, circularly shifted to a common center (0°, by convention), and averaged, yielding a single reconstruction per participant, time point, cue condition (i.e., informative vs. uninformative) and task (i.e., pro vs. anti-cue). Condition-wise channel response functions were averaged, converted to polar form, and projected onto a vector with angle 0°:

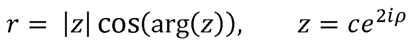

Where *c* is a vector of estimate channel responses and ρ is a vector of angles at which the channels peak. To ensure internal reliability, this entire analysis was repeated 100 times, and unique (randomly chosen) subsets of trials were used to define the training and test data sets during each iteration. The results were then averaged across permutations to obtain a single reconstruction strength estimate for each participant, task condition, and timepoint.

### Statistical Comparisons – Behavioral Data

Participants’ behavioral data (i.e., absolute average recall error and average response time; Figure 1B-E) were analyzed using standard repeated-measures parametric statistics (e.g., t-test, ANOVA); for these comparisons we report test statistics, p-values, and effect size estimates.

### Statistical Comparisons – N2pc, Decoding Performance, and Inverted Encoding Model

The decoding analysis and inverted encoding model we used assume chance-level performance of 0. Likewise, direct comparisons of decoding performance or reconstruction strength across conditions (e.g., pro-cue vs. anti-cue) assume null statistics of 0. Thus, we evaluated task-relevant and task-irrelevant decoding performance by generating null distributions of decoding performance (or differences in decoding performance across conditions) by randomly inverting the sign of each participant’s data with 50% probability and averaging the data across participants. This procedure was repeated 10,000 times, yielding a 10,000-element null distribution for each time point. Finally, we implemented a cluster-based permutation test (Maris & Oostenveld, 2007) with cluster-forming and cluster-size thresholds of p < 0.05 to correct for multiple comparisons across time points.

## Results

We recorded EEG while 40 human volunteers performed a retrospectively cued spatial recall task (Figure 1A). Participants remembered the positions of two discs over a brief delay, and a retrospective color cue presented 1.25 seconds later instructed participants to continue remembering the positions of both discs (i.e., uninformative trials) or to prioritize one of the discs for subsequent recall (i.e., informative trials). The locations of the two discs were fully randomized across experimental blocks (subject to the constraint that two discs could not appear at the same location on a given trial). At the end of the trial, participants recalled the position of the task-relevant disc via mouse click. Behavioral performance was quantified via average response times and average absolute recall error (i.e., the average absolute difference between the correct and reported disc position). In separate experimental blocks, participants performed a pro-cue task or an anti-cue task. During the pro-cue task informative cues were assigned 100% validity; during the anti-cue task informative cues were assigned 0% validity (i.e., the cue color indicated which disc was task-irrelevant). This allowed us to disentangle the effects of endogenous and exogenous factors on the selection of WM content: during the pro-cue task the color cue indicates which of the two remembered stimuli are task relevant, and endogenous and exogenous selection mechanisms are aligned. During the anti-cue task, however, the color cue indicates which of the two stimuli are task-irrelevant, placing endogenous and exogenous selection mechanisms in competition (e.g., van Ede et al., 2020). Task order (i.e., pro-followed by anti-cue or vice versa) was counterbalanced across participants, and participants were explicitly reminded about cue validity at the beginning of every block.

### Endogenous and Exogenous Factors Influence the Selection of Task-Relevant WM Content, but in Different Ways

A two-factor repeated measures analysis of variance (ANOVA) applied to participants’ average absolute recall errors (Fig 1B) revealed a main effect of cue type (i.e., informative vs. uninformative; [F(1,39) = 15.854, p = 0.0003, η = 0.289]), with lower errors during informative vs. uninformative cue trials. Likewise, this analysis revealed a main effect of task (i.e., pro- vs. anti-cues; [F(1,39) = 8.168, p = 0.0068, η = 0.1732]), with lower errors during the pro- vs. anti-cue task, and a significant interaction between these factors [F(1,39) = 5.35, p = 0.0261]. A complementary analysis of response times (Fig 1D) revealed a main effect of cue type [F(1,39) = 483.046, p < 0.0001, η = 0.925], with response times faster during informative vs. uninformative cue trials, no main effect of task [F(1,39) = 0.022, p = 0.884, η = 0.060], and a significant interaction between these factors [F(1,39) = 30.362, p < 0.0001].

In planned comparisons, we sought further clarity on how endogenous and exogenous factors influenced participants’ memory performance. We reasoned that during informative pro-cue trials endogenous and exogenous factors are aligned while during informative anti-cue trials they conflict. Thus, to isolate the effects of exogenous factors on memory performance we compared participants’ recall errors and response times across informative pro- and anti-cue trials (i.e., the simple effect of task for informative cues). Conversely, to quantify the effect of endogenous selection on memory performance, we first calculated differences in participants’ recall errors and response times across informative and uninformative trials in the pro-cue task (i.e., the simple effect of cue type for the pro-cue task). Next, we subtracted the effects of exogenous factors estimated in the previous step from these differences to isolate the effects of endogenous factors on memory performance (i.e., while accounting for the fact that during the pro-cue task endogenous and exogenous cues are aligned while during the anti-cue task they are opposed). Endogenous factors had a facilitatory effect on task performance, lowering recall errors (M = 1.78°; 95% CI = 0.645°-3.112°; Fig 1C) and speeding response times (M = 0.165 sec; 95% CI = 0.014-0.305 sec; Fig 1E). In contrast, exogenous factors significantly worsened participants’ recall errors (M = 0.961°; 95% CI = 0.191°-1.863°; Fig 1C) but had no effect on response times (M = - 0.055, 95% CI = -0.073-0.189; Fig 1E). Thus, endogenous and exogenous factors had faciliatory and deleterious effects on participants’ memory performance, respectively.

### Manipulation Check: The Anti-cue Task Requires a Greater Degree of Cognitive Control than the Pro-cue task

A key assumption of our experimental approach holds that the anti-cue task produces conflict between endogenous and exogenous selection mechanisms. We reasoned that cognitive control is needed to resolve this competition and drive the selection of task-relevant WM content, and that therefore a greater degree of cognitive control would be required during the anti-cue task compared to the pro-cue task (i.e., when endogenous and exogenous selection mechanisms are aligned). We tested this prediction by estimating and comparing theta power (4-7 Hz) over frontal electrode sites during the pro- and anti-cue tasks. Frontal theta power has robustly linked with the need for cognitive control (Cavanagh & Frank, 2014), scales with WM load (Jensen & Tesche, 2002), and predicts successful working memory updating (Itthipurupat et al., 2013). Thus, we expected larger frontal theta power estimates during the anti-cue vs. the pro-cue task. Indeed, we observed significantly greater frontal theta power during the anti-cue vs. pro-cue task that was maximal over frontal midline electrode sites (Figure 2). Note that this effect emerged only after presentation of the retrocue, consistent with a need for “online” cognitive control rather than a general increase in difficulty during the anti-cue vs. pro-cue task. These data support our contention that the anti-cue task produces significant conflict between endogenous and exogenous selection mechanisms.

### Competition Between Endogenous and Exogenous Selection Delays the Selection of Task-relevant WM Content

To understand how competition between endogenous and exogenous factors influence the selection of WM content, we examined how pro- and anti-cues influenced our ability to decode stimulus positions from scalp EEG. Our approach builds on studies demonstrating that stimulus- and location-specific information can be decoded from alpha-band EEG signals (e.g., Foster et al., 2016), and that attending to an item or location stored in WM selectively boosts decoding for the attended information (e.g., Lewis-Peacock et al., 2012; LaRocque et al., 2013; Sprague et al., 2016; Ester et al., 2018). We implemented a multivariate distance-based decoding analysis (Wolff et al., 2017) that was customized for our (parametric, circular) location space. This approach is similar to image reconstruction techniques (i.e., “inverted encoding models”) but does not require the experimenter to specify a specific coding model or basis set. To facilitate comparisons across cue conditions and tasks, participant-level decoding time series were sorted by task relevance: during the pro-cue task decoding performance for the cue-matching disc was designated task-relevant and decoding performance for the cue-nonmatching disc was designated task-irrelevant; during the anti-cue task this mapping was reversed.

We tested two models describing how competition between endogenous and exogenous selection mechanisms influences the prioritization of task-relevant and task-irrelevant WM content. The first model – which we term “retro-capture” – was motivated by studies reporting exogenous shifts of attention to task-irrelevant stimuli in the external attention literature (e.g., Theeuwes, 1992; Folk et al., 1992). This model predicts a transient increase in position decoding performance for the cue-matching but task-irrelevant stimulus during the anti-cue task, followed by a later increase in position decoding performance for the cue-nonmatching but task-relevant position (i.e., after the effects of selecting the task-irrelevant stimulus have been resolved). The second model – which we term “delayed selection” - predicts that competition between endogenous and exogenous selection mechanisms merely delays the selection of task-relevant WM content until this competition is resolved. Thus, this model predicts a significant delay in the onset of above-chance position decoding for the cue-nonmatching but task-irrelevant stimulus anti- vs. pro-cue task, but no evidence for above-chance decoding of the cue-matching but task-irrelevant stimulus during the anti-cue task.

Our experimental task (Figure 1A) was deliberately constructed so that the effects of endogenous and exogenous factors on the selection of WM contents could be measured during informative *and* uninformative trials. For example, during uninformative trials participants received an uninformative retrospective cue instructing them to remember the positions of both discs. Upon presentation of the probe display, this uninformative cue was replaced by a 100% valid (pro-) or 0% valid (anti-) cue instructing participants which disc to report. Conversely, during informative trials pro- and anti-cues were presented midway through the storage period. Since informative and uninformative trials had different response demands (i.e., pro- and anti-cues presented at the end of uninformative trials required an immediate response while pro- and anti-cues presented during the memory delay during informative trials did not), we analyzed data from these conditions separately.

We first considered data from uninformative cue trials (Figure 3). Task-relevant and task-irrelevant location decoding performance in the pro-cue (Figure 3A) and anti-cue (Figyre 3B) tasks increased rapidly during the sample display but returned to chance levels by the time the (uninformative) retrocue was presented 1.75 sec later. Task-irrelevant decoding performance remained at chance levels through the retrocue and probe displays while task-relevant decoding performance increased from chance-to above-chance levels during the probe period. Visual comparisons of probe-locked task-relevant decoding performance suggested that above-chance decoding performance was reached earlier during the pro-relative to the anti-cue task (Figure 3C). To quantify this effect, we extracted and compared probe-locked task-relevant decoding time courses during the pro- and anti-cue tasks via cross-correlation. Specifically, we computed correlations between the timeseries of task-relevant decoding performance during the pro- and anti-cue tasks while temporally shifting the former by - 1.0 to +1.0 sec in 4 msec intervals relative to the latter, yielding a correlation-by-lag function (see Methods). Observed cross-correlation coefficients (Figure 3D) exceeded those expected by chance over lags spanning -0.33 to -0.22 sec and fell below those expected by chance over a period spanning +0.25 to +0.35 sec, confirming that task-relevant decoding performance reached above chance levels earlier during the pro- vs. anti-cue task.

A complementary analysis of cue-locked decoding performance during informative trials yielded a nearly identical pattern of findings (Figure 4). Specifically, we once again found no evidence for above-chance decoding of the cue-matching but task-irrelevant stimulus position during the anti-cue task (Figure 4B). We did, however, observe a significant delay in the onset of task-relevant decoding performance during the anti- vs. pro-cue tasks (Figure 4C-D). Thus, the results of probe- and cue-display-locked position decoding performance reveal (a) no evidence for above-chance decoding of the cue- or probe-matching but task-irrelevant stimulus position (Figure 3B & 4B) and (b) a significant delay in the onset of above-chance decoding of the task-relevant stimulus position compared to the pro-cue task (Figures 3C-D & 4 C-D). These findings are incompatible with a model of internal selective attention where competition between exogenous and endogenous selection mechanisms can produce inadvertent shifts of attention to cue-matching but task-irrelevant WM content.

**Figure 4.**
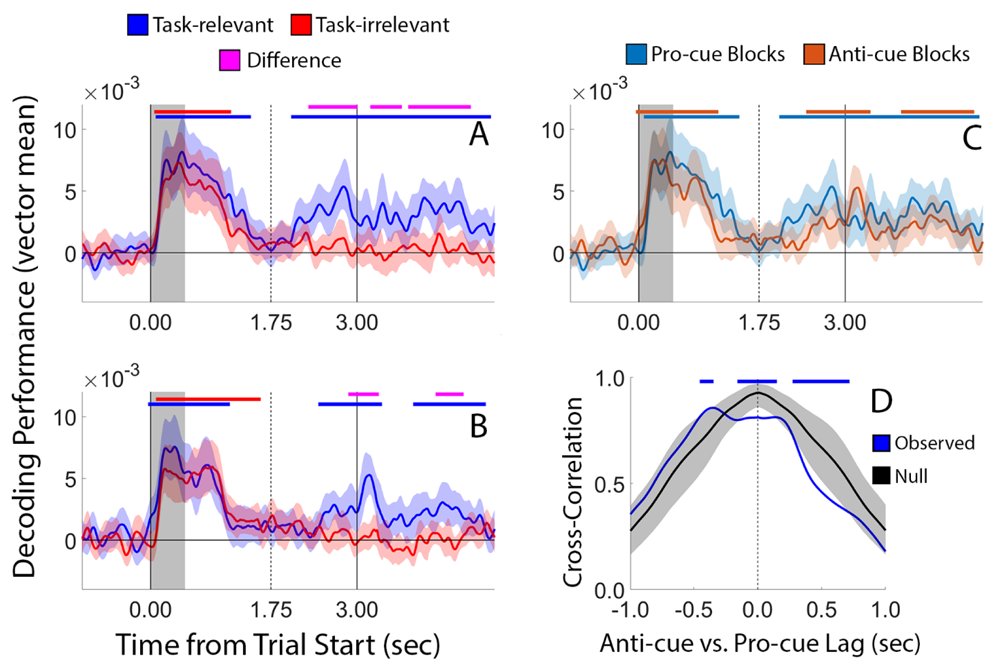
Location Decoding Performance During Informative Trials. (A, B) Decoding performance for task-relevant and task-irrelevant locations during pro-cue and anti-cue blocks, respectively. (C) Overlay of task-relevant location decoding performance for pro-cue and anti-cue blocks (i.e., the blue lines in panels A and B). (D) Cross-correlation analysis showing a significant delay in the onset of above-chance probe-locked task-relevant decoding performance during the anti- vs. pro-cue task. All conventions are identical to Figure 3.

A motivated critic could dismiss our conclusions as based on a null result. For example, perhaps our anti-cue task was insufficient to produce selection of cue-matching yet task-irrelevant stimuli. This argument is difficult to reconcile with behavioral findings showing clear memory impairments during the anti- vs. pro-cue task (Figure 1C) and higher cue-locked frontal theta power during the anti- vs. pro-cue task (Figure 2) uninformative and informative cue trials in the anti- vs. pro-cue tasks (Figure 2). A second possibility is that the parametric similarity-based decoding approach we used is somehow insensitive to resolve the selection of cue-matching but task-irrelevant WM content during the anti-cue task. We tested this possibility by re-analyzing data from informative cue trials using a support vector machine (SVM) based decoding approach (Figure S5) and an inverted encoding model (Figure 6). SVM-based decoding failed to reveal above-chance decoding of the cue-matching but task-irrelevant position during WM trials (Figure 5B), along with a significant delay in task-relevant decoding during the anti- vs. pro-cue task (Figure 5C). Likewise, the results of the inverted encoding model analysis are a perfect qualitative replication of the pattern reported in Figure 4: we observed no evidence for robust above-chance representations of the cue-matching but task-irrelevant stimulus during the anti-cue task (Figure 6B) and a significant delay in above-chance reconstructions for the task-relevant position during the anti- vs. the pro-cue task (Figure 6C). Thus, there is little reason to suspect that our key findings (Figures 3-4) are due to the specific decoding approach we used.

**Figure 5.**
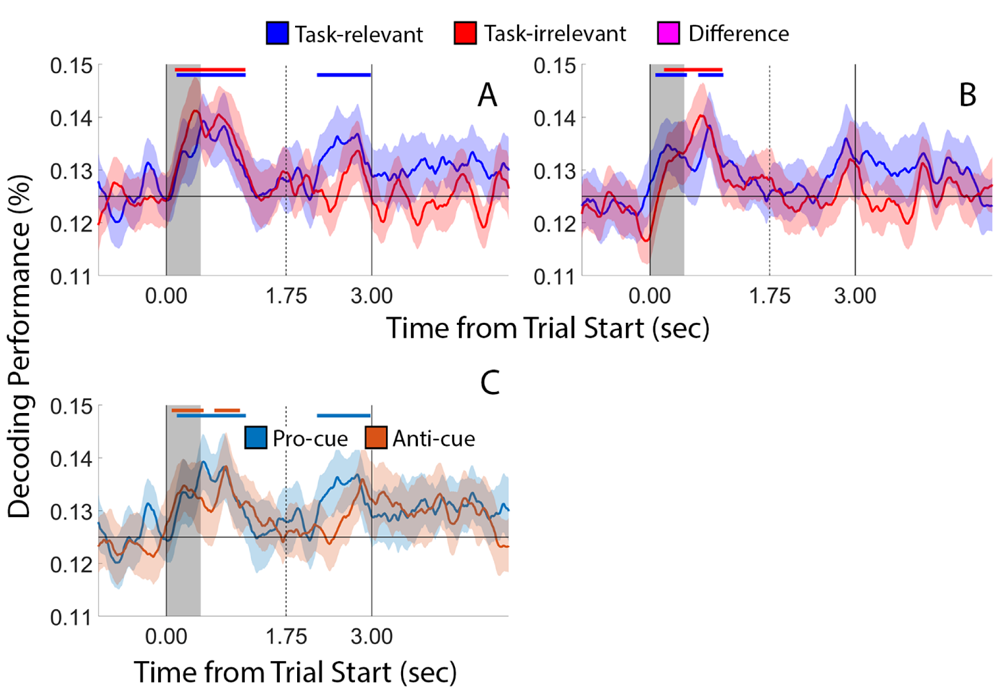
Support Vector Machine-based Decoding of Stimulus Position. To ensure the generality of our findings (e.g., Figure 4), we decoded the positions of the task-relevant and - irrelevant discs during the pro-cue task (A) and the anti-cue task (B). Plotting conventions are identical to Figure 4. We did not perform a cross-correlation analysis due to the absence of above-chance decoding of the cue-matching but task-irrelevant stimulus during the anti-cue task.

**Figure 6.**
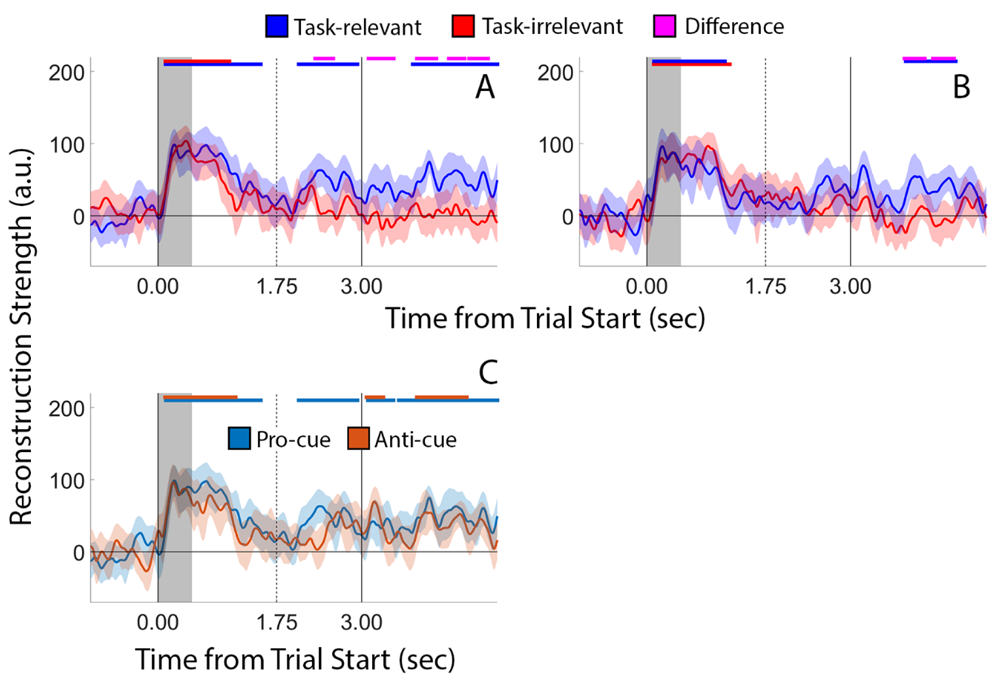
Inverted Encoding Model Analysis. We modeled patterns of alpha-band activity at each electrode site as a weighted combination of eight position filters, each with an idealized tuning curve. Filter weights from each electrode were used to reconstruct a representation of remembered position(s) in an independent test data set. Conventions are identical to Figure 5.

Next, we considered the possibility that our decoding approach (Figures 3-4) lacked the temporal sensitivity to detect the selection of task-irrelevant WM content. For example, perhaps the temporal dynamics of changes in alpha power are insufficient to measure weak or intermittent (i.e., occurring on only a subset of trials) shifts of attention to the cue-matching but task-irrelevant position during the anti-cue task. We investigated this possibility by tracking the N2pc, an event-related potential (ERP) component known to track covert shifts of attention across visual hemifields. The N2pc is a difference wave defined by greater negative voltages over occipitoparietal electrode sites contralateral (vs. ipsilateral) to a visual target beginning ∼200 ms after stimulus onset (Luck & Hillyard, 1994) and can be used to track endogenously and exogenously driven shifts of covert attention with exceptionally high temporal precision (Hickey et al., 2006; Burra & Kerzel, 2013). We reasoned that if competition between endogenous and exogenous selection mechanisms produces a selection of cue-matching but task-irrelevant information, then we should observe a significant N2pc over electrode sites contralateral to the visual hemifield containing the cue-matching but task-irrelevant disc during the anti-cue task. Conversely, if competition between endogenous and exogenous selection mechanisms delays the selection of task-relevant WM content, then we should (a) observe a robust N2pc over electrode sites contralateral to the visual hemifield containing the task-relevant disc during the pro-cue task, and (b) observe a significant delay in the onset of the N2pc over electrode sites contralateral to the visual hemifield containing the task-relevant disc during the anti-cue vs. pro-cue task.

To test these predictions, we computed voltage differences over occipitoparietal electrode sites contralateral to the visual hemifield containing the task-relevant disc during the pro- and anti-cue tasks. To control for possible sensory confounds we restricted our analyses to trials where the task-relevant and task-irrelevant discs appeared in opposite visual hemifields (approximately 70 trials/task). The N2pc was defined as the average voltage difference over contralateral and ipsilateral electrodes spanning 200-300 ms after cue onset. Since we defined the N2pc with respect to the visual hemifield containing the task-relevant disc, and since we restricted our analysis to trials where the task-relevant and task-irrelevant discs appeared in opposite visual hemifields, shifts of attention towards the cue-matching but task-irrelevant disc during the anti-cue task should manifest as a positive-going waveform 200-300 ms after cue onset.

We observed a statistically robust N2pc from 200-300 ms following the appearance of an informative pro-cue (Figure 7), indicating that participants executed a shift of covert visual attention to the visual hemifield containing the cue-matching and task-relevant disc. Conversely, we observed no evidence for a positive-going waveform during the same interval following the presentation of an informative anti-cue. That is, we found no evidence suggesting that participants executed a shift of covert spatial attention towards the visual hemifield containing the cue-matching but task-irrelevant disc during anti-cue trials. Instead, we observed a robust negative-going difference wave beginning ∼350 ms after the appearance of an anti-cue. We speculate that this negative-going difference wave is identical to the N2pc elicited during the pro-cue task whose onset has been delayed by competition between endogenous and exogenous selection mechanisms. Nevertheless, the results of this analysis provide converging evidence against the hypothesis that competition between endogenous and exogenous selection mechanisms drives the inadvertent selection of cue-matching but task-irrelevant information.

**Figure 7.**
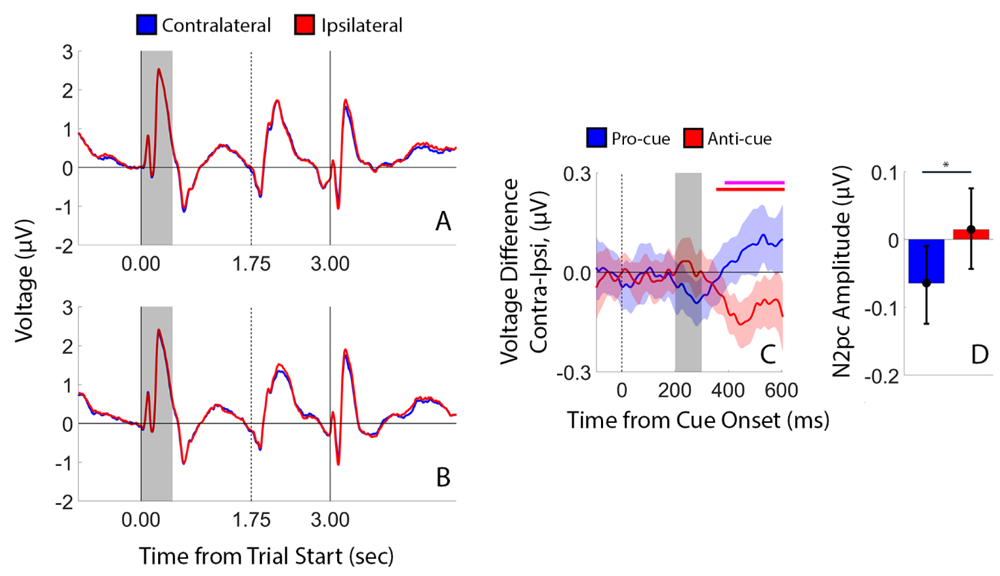
Event-related Potentials Reveal Delayed Selection of Task-relevant WM Content During the Anti-Cue Task. (A) Average contralateral and ipsilateral ERP waveforms during the pro-cue task, time-locked to trial start (0.00 sec). The vertical lines at times 1.75 and 3.00 sec depict the onset of the retrocue and probe displays, respectively. The shaded region depicts the duration of the sample display. (B) Identical to (A), but for the anti-cue task. (C). Difference waves (i.e., contralateral-ipsilateral) time locked to retrocue onset (time 0 ms). The shaded region spanning 200-300 ms depicts the canonical N2pc window. Horizontal bars at the top of the plot mark epochs where difference wave voltage was significantly greater than chance (red bar) or when anti-cue difference wave voltage was significantly greater than pro-cue difference wave voltage (maroon). Shaded regions depict the 95% confidence interval of the mean. (D) N2pc amplitudes, defined as the average difference wave voltage over a period spanning 200-300 ms after cue onset. Error bars depict the 95% confidence interval of the mean; *, p< 0.05, bootstrap test.

### Other Alternative Explanations

Next, we considered the possibility that evidence for the selection of the task-irrelevant disc during the anti-cue task was obscured by trial averaging. For example, perhaps the selection effect is small, short, lived, or intermittent (i.e., occurring on only a subset of trials). We tested this possibility by recomputing alpha-band-based decoding performance for the task-irrelevant disc after sorting participants’ anti-cue task performance by median recall error (i.e., “high” vs. “low”). Here, we reasoned that since exogenous factors have a deleterious effect on participants’ recall errors during the anti-cue task (Figure 1C), exogenous selection of the task-irrelevant disc – as indexed by higher task-irrelevant decoding performance – should be more evident during high recall error trials. However, this was not the case: we observed no evidence for above-chance task-irrelevant decoding performance during low-or high-error informative (Figure 8A) or uninformative trials (Figure 8B). Thus, it is unlikely that the pattern of exogenous-then-endogenous selection predicted by the retro-capture model was obscured by trial-averaging.

**Figure 8.**
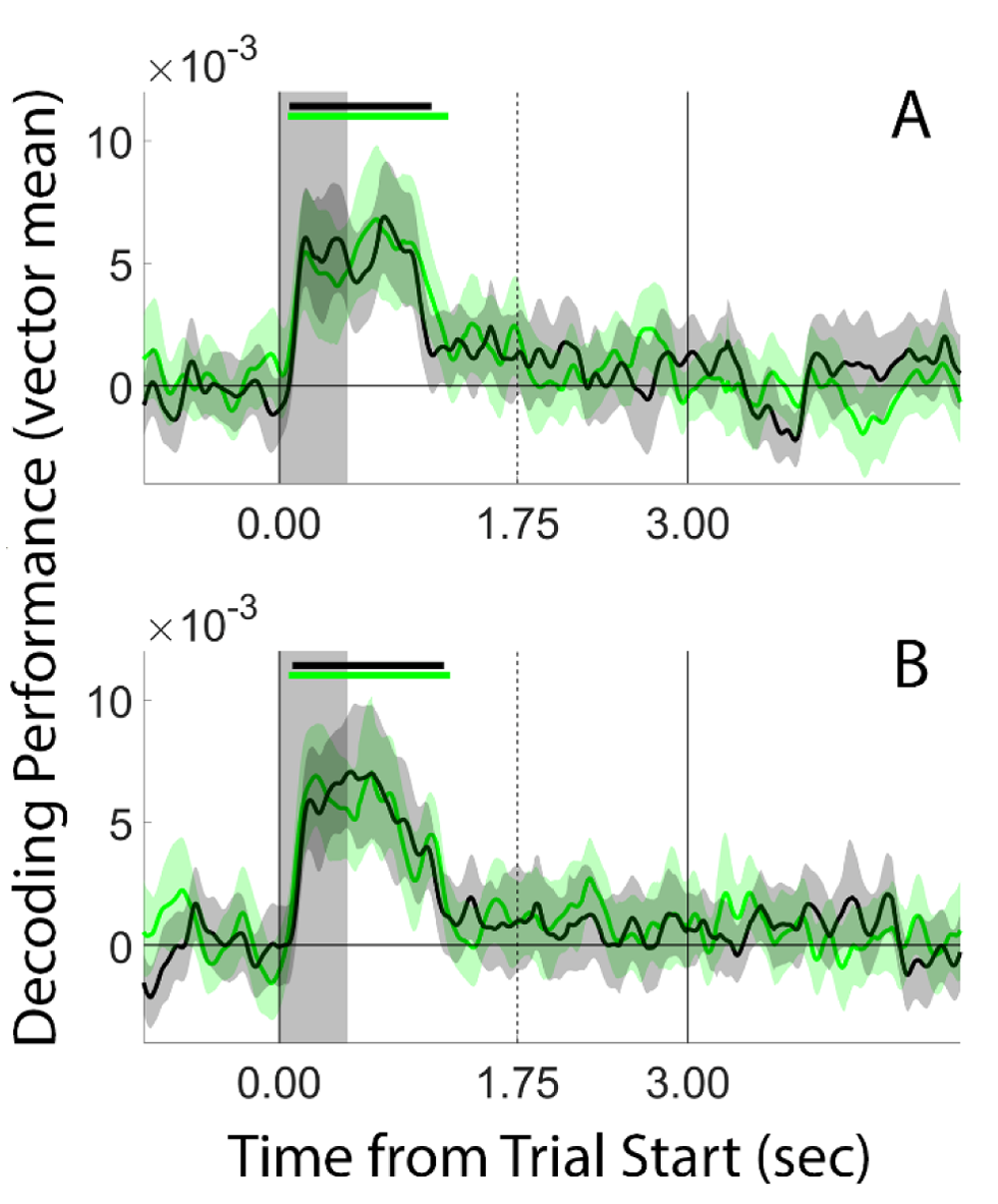
Split-half Analysis of Task-irrelevant Decoding Performance During the Anti-cue Task. To examine whether exogenous selection of the task-irrelevant disc during anti-cue blocks was obscured by trial averaging, we sorted task-irrelevant decoding performance during neutral (A) and informative (B) trials by participants’ recall errors. We reasoned that since exogenous factors have a deleterious effect on participants’ recall errors (Fig 1C), exogenous selection of the task-irrelevant disc – as indexed by higher task-irrelevant decoding performance – should be more evident during high recall error trials (black lines) than low recall error trials (green lines). However, we observed no evidence for above-chance task-irrelevant decoding performance in any of the conditions we examined. Plotting conventions are identical to those in Figure 4.

We also considered the hypothesis that selection of the cue-matching but task-irrelevant disc during the anti-cue task was obscured by successful cognitive control. Specifically, we reasoned that shifts of attention towards the location of the task-irrelevant disc might be more likely during trials contaminated by lapses of attention. To test this hypothesis, we re-computed cue-matching but task-irrelevant location decoding performance after sorting participants’ alpha-band EEG data by frontal theta power (Figure 2), reasoning that inadvertent selection of cue-matching but task-irrelevant stimuli would be more likely during trials where frontal theta power (indexing cognitive control) was low vs. high. However, we observed no evidence for above-chance decoding of the cue-matching but task-irrelevant position during high-or low-theta power trials (Figure 9). This analysis provides converging evidence suggesting that exogenous factors do not lead to a selection or re-activation of cue-matching but task-irrelevant WM content, but instead delay the endogenous selection of task-relevant WM content.

**Figure 9.**
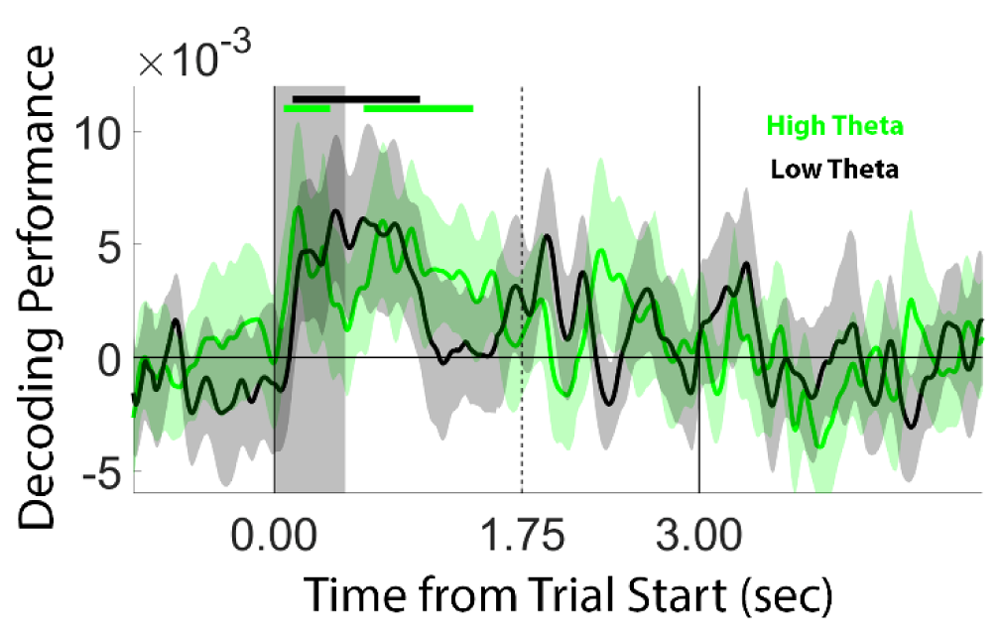
Task-irrelevant Decoding Performance During the Anti-Cue Task Sorted by Frontal Theta Power. Conventions are identical to Figure 4B.

Finally, although our analyses reveal no evidence for a selection of cue-matching but task-irrelevant information during the anti-cue task, they do reveal a significant delay in the selection of task-relevant information during the anti- vs. pro-cue tasks (Figures 3C-D and 4C-D). This effect could reflect a delay in the selection of task-relevant information caused by competition between endogenous and exogenous selection mechanisms during the anti-cue task or some other task-specific factor. For example, one trivial possibility is that it simply takes participants longer to interpret anti-cues vs. pro-cues. However, this explanation is difficult to reconcile with the fact that neither the main effect of task (i.e., pro-cue vs. anti-cue; Figure 1D) nor the simple effect of task (Figure 1E) on response times during informative cue trials reached significance. A second possibility is that delayed above-chance decoding performance during the anti-cue task was caused by carryover effects. For example, although task order was counterbalanced across observers, perhaps participants who completed the pro-cue task followed by the anti-cue task had extra difficulty interpreting anti-cues compared to participants who performed the anti-cue task followed by the pro-cue task. To test this possibility, we compared the time-courses of task-relevant decoding performance during informative anti-cue trials in participants who performed the pro-cue task followed by the anti-cue task (N = 17) or vice versa (N = 23). For both groups, task-relevant decoding performance reached above chance levels shortly before or immediately after the onset of the probe display (Figure 10). If anything, the onset of above-chance decoding performance occurred later for participants who performed the anti-cue task second vs. those who performed the anti-cue task first, though this difference was not significant (p = 0.146; randomization test, see Methods). Thus, order effects cannot account for delays in task-relevant decoding performance during the anti-cue vs. pro-cue blocks.

**Figure 10.**
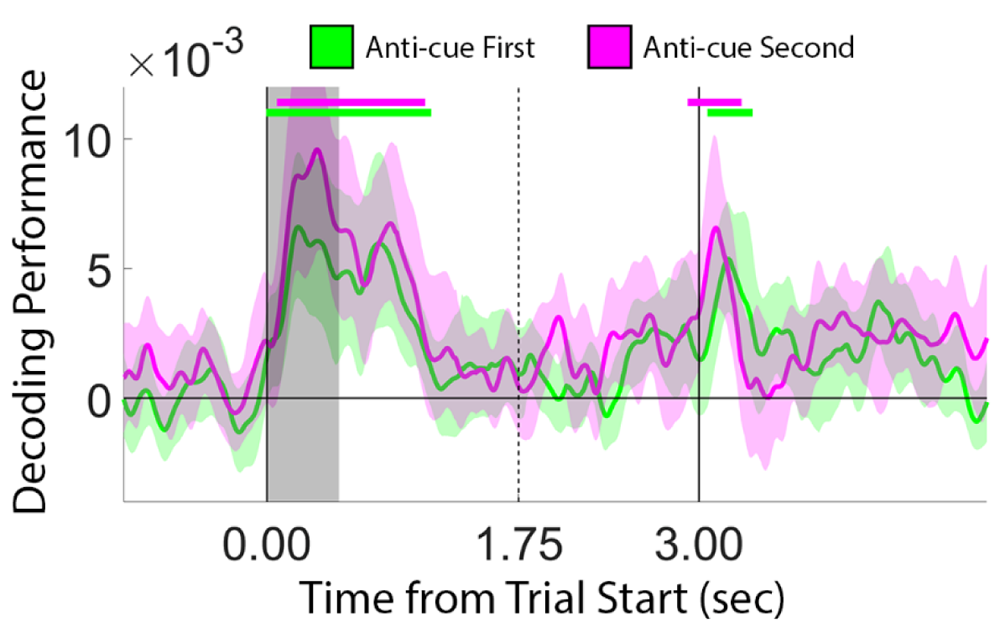
Delayed Improvements in Task-Relevant Decoding Performance During the Anti-Cue Task Cannot be Explained by Order Effects. We tested whether delayed improvements in task-relevant decoding performance during the anti-cue (vs. pro-cue) task were caused by order effects by splitting decoding performance across participants who performed the anticue task followed by the pro-cue task (green) or vice versa (maroon). If anything, above-chance decoding performance was reached earlier for participants who completed the pro-cue followed by the anti-cue tasks, though this effect was not significant (p = 0.141; randomization test).

## Discussion

Selective attention can be allocated to sensory inputs and internal representations based on voluntary, endogenous factors or involuntary, exogenous factors. An enormous literature suggests that external selection is mediated by competition between endogenous and exogenous factors, with the focus of selection determined by the balance of competition between these factors (e.g., Wolfe, 2020). Endogenous and exogenous factors may also compete to control the selection of internal representations, for example, those stored in WM (e.g., van Ede et al., 2020). Here, we show that - unlike external attention – this competition does not result in a selection of task-irrelevant stimuli. This, in turn, supports the hypothesis that internal and external selective attention are mediated by at least partially non-overlapping mechanisms.

A motivated critic could dismiss our conclusion as based on a null result. While it is certainly true that absence of evidence for the selection of task-irrelevant stimuli during the anti-cue task need not imply evidence against such an effect, our conclusion is based on a series of control analyses that systematically exclude multiple explanations for why such an effect did not appear. First, perhaps our experimental approach was insufficient at creating conditions conducive to the exogenous selection of cue-matching but task-irrelevant stimuli during the anti-cue task. While we cannot fully exclude this possibility, we note that participants’ memory performance was significantly worse during the anti- vs. pro-cue tasks (Figure 1B) and that the appearance of an anti-cue led to a significant increase in frontal theta power compared to the appearance of a pro-cue (Figure 2), consistent with a need for greater cognitive control during the anti- vs. pro-cue task. Second, perhaps our specific decoding approach lacked the sensitivity to identify the selection of task-irrelevant information during the anti-cue task. Again, it is difficult to fully exclude this possibility, but we note that we observed qualitatively different patterns of findings across two different decoding methods (similarity-based vs. support vector machine-based; Figures 4 and 5, respectively) and the results of an inverted encoding model analysis where we reconstructed remembered positions from EEG activity (Figure 6). Third, perhaps the posterior alpha-band signal (8-13 Hz) lacks the temporal resolution necessary to resolve fleeting or intermittent selection of task-irrelevant information during the anti-cue task. However, analyses of the N2pc ERP component responses (with a temporal resolution of ∼4 ms) revealed clear evidence for the selection of the task-relevant disc during the pro- and anti-cue tasks but no evidence for the selection of the task-irrelevant disc during the anti-cue task (Figure 7). Fourth, a variety of additional control analyses demonstrate that the selection of task-irrelevant information during the anti-cue task was not obscured by high behavioral performance (Figure 8), successful cognitive control (Figure 9), or task order effects (Figure 10). Importantly, in many of these analyses we *did* find evidence for a temporal delay in the selection of task-relevant WM content during the anti-cue vs. the pro-cue task. Thus, we argue that unlike external attention – where competition between endogenous and exogenous selection mechanisms produces clear evidence for the selection of irrelevant stimuli – competition between endogenous and endogenous internal selection mechanisms does not produce a selection of task-irrelevant memory content and is resolved in a fundamentally different way. More generally, this result points towards important differences in how voluntary and involuntary selection mechanisms compete to control the processing of external sensory inputs vs. internal memory representations.

The current findings may inform neurocomputational models of WM. For example, conjunctive coding models predict that WM representations are maintained by spiking activity in feature- and/or location-specific neural populations (e.g., Schneegans & Bays, 2017; 2018). While the exact mechanisms vary by implementation, these models generally predict that a feature probe in one dimension (e.g., orientation) activates spiking patterns in neural populations that code this feature and those that code other features of the same object (e.g., color) and/or its location. This, in turn, enables robust read-out of the probed and non-probed stimulus dimensions by downstream neural populations. While these models were not developed to describe the anti-cue task contemplated here, one could reasonably predict an increase in task-irrelevant decoding performance after presentation of an anti-cue based on their general architecture. We observed no evidence for such an effect, and it remains to be seen whether these models can be modified to predict behavioral and neural data during pro- and anti-cue tasks. Alternately, pattern completion models predict that the contents of WM reside in different neural states – an “active” state mediated by sustained spiking activity and a “latent” state mediated by short-term synaptic plasticity (e.g., Mongillo et al., 2008; Manohar et al., 2019). Presentation of a feature probe that matches a stimulus stored in a latent format reinstates activity patterns evoked when that stimulus was encoding, prompting and/or “refreshing” of the neural representation of the probe-matching item through pattern completion. This prediction enjoys some support: a representation stored in WM item can be “re-activated” (as indexed by above-chance EEG decoding performance) by a task-irrelevant sensory input (Wolff et al., 2017) or a transcranial magnetic stimulation (TMS) pulse applied over sensory cortex (Rose et al., 2016). Conversely, in the current study we found no evidence for a reactivation of cue-matching but task-irrelevant WM content following presentation of an anti-cue. However, one salient difference between the current study and prior work is that in the latter, an informative retrospective cue instructed presented prior to the “impulse” stimulus instructed participants which of two remembered stimuli should be prioritized for report. Thus, re-activation of information stored in synaptic traces may be contingent on the network responsible for storing information to be selected or otherwise primed for decision making and action.

A recent study reported that the onset of small-but-robust biases in gaze position (referred to hereafter as microsaccades) towards the memorized location of a task-relevant stimulus were delayed following the appearance of an anti- vs. pro-cue (van Ede et al., 2020). The authors of this study speculated that delays in selecting task-relevant information during an anti-cue task were caused by an automatic “refreshing” of cue-matching but task-irrelevant WM content. However, as the authors of this study acknowledged, this hypothesis cannot be substantiated based on microsaccades alone. First, although attention-related modulations of cortical and subcortical processing are larger during trials containing microsaccades towards the location of a (covertly) attended stimulus, clear attention-related modulations are also observed in the absence of microsaccades (Yu et al., 2022; Liu et al., 2022; though see Lowet et al., 2018 for a contradictory result). Second, although we did not (and could not, lacking a functional eyetracker at the time this study was conducted) record eye position with sufficient spatiotemporal resolution needed to replicate the findings reported by (van Ede et al., 2020), we observed no evidence for an enhancement of cue-matching but task-irrelevant position information during the anti-cue task that would be predicted by the account endorsed by van Ede and colleagues (2020). Thus, we speculate that the delays in microsaccade biases reported by these authors were generated by delayed selection of the appropriate WM content. Further research could compare the time-courses of oculomotor and neural signals associated with internal selection to test this possibility.

To summarize, the current findings support recent suggestions that endogenous and exogenous selection mechanisms compete to control access to internal WM representations. However, this competition is resolved in a fundamentally different way than that seen during external attention. Specifically, endogenous and exogenous competition does not produce an errant selection or refreshing of salient but task-irrelevant WM content. Instead, this competition delays the selection of task-relevant memory content by endogenous mechanisms. This result reveals a fundamental difference between the operation of external and internal attentional selection.

